# Myelin-Free Nuclei Isolation from Mouse Hippocampus and Cerebellum for snRNA-Seq with Benchtop Gradient Centrifugation

**DOI:** 10.64898/2026.04.03.716374

**Authors:** Benu George, Braedon Q Kirkpatrick, Victor Kilonzo, Qiang Zhang

**Author notes:** **Correspondence to**, 169 Newton Road Department of Neurology, Roy J. and Lucille A. Carver College of Medicine University of Iowa, Iowa City, IA 52242.

## Abstract

Nuclei isolation from myelin-rich adult mouse brain regions remains challenging for single-nucleus RNA sequencing because myelin and debris can reduce nuclei quality. We describe an optimized protocol for mouse hippocampi and cerebella using tube-and-pestle homogenization, low-volume sucrose-gradient pelleting with a standard benchtop centrifuge, and magnetic enrichment as the recommended final cleanup step to reduce debris/non-nuclear carryover. Under the tested conditions, the workflow produces intact, debris-reduced nuclei and supports downstream 10x Genomics Flex and PARSE WT library preparation.

**Graphical abstract:** 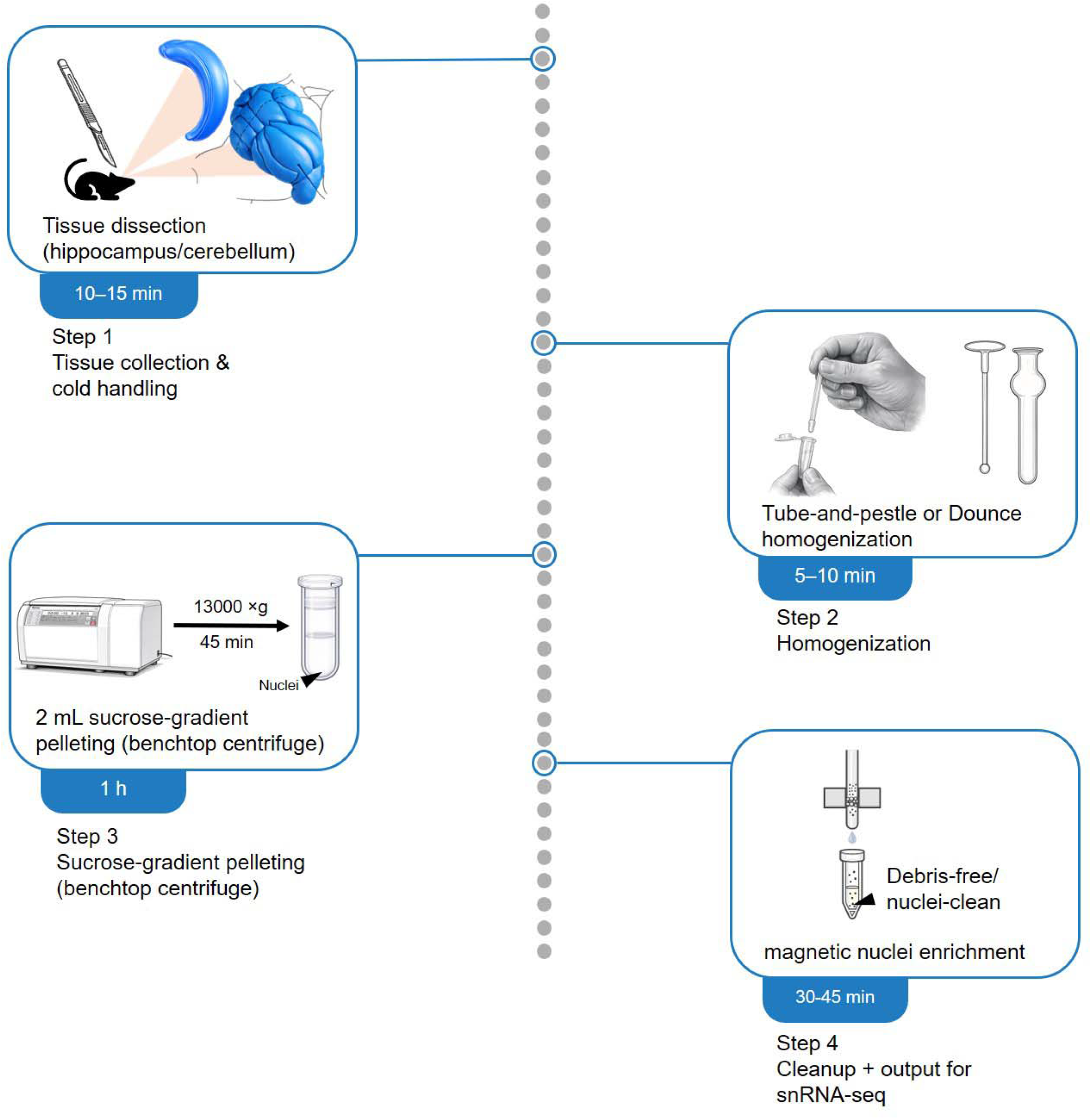

**Highlights:**

- Benchtop sucrose-gradient pelleting enables rapid nuclei purification from myelin-rich adult mouse brain
- Scales across tissue inputs (e.g., hippocampus ∼15–20 mg; cerebellum ∼50–70 mg) without ultracentrifugation or 15 mL gradients
- Magnetic enrichment as the recommended final cleanup step further reduces myelin/debris carryover and is compatible with 10x Flex and PARSE WT workflows.

## BEFORE YOU BEGIN

Single-nucleus RNA sequencing (snRNA-seq) requires isolation of intact nuclei with low debris and myelin carryover to support downstream library preparation and sequencing quality (1,2). This protocol describes nuclei isolation from adult mouse hippocampi and cerebella for downstream snRNA-seq using 10x Genomics Flex and PARSE WT workflows. It is optimized for myelin-rich adult mouse brain tissue, where residual myelin and fine debris can complicate nuclei cleanup and downstream handling (1–3). Although the procedure below describes hippocampus- and cerebellum-derived samples, it may be adapted to other adult mouse brain regions with re-optimization of tissue input, homogenization intensity, and cleanup conditions.

Before starting, prepare all reagents and equipment in advance, keep samples and buffers on ice throughout processing, and organize the workflow to minimize delays between homogenization, gradient pelleting, and cleanup.

1. **Prepare the work area and temperature-controlled equipment**

a. Pre-cool the benchtop centrifuge(s) and rotor(s) used for nuclei isolation steps to 4°C.
b. Place tube racks, pestles, and centrifuge-compatible tubes on ice.
c. Arrange the workspace so reagents and tubes are positioned in order of use to support continuous processing.
2. **Prepare buffers and cleanup reagents**

a. Prepare nuclei isolation, wash/resuspension, and cleanup buffers as described in the Materials and Equipment section.
b. Prepare the sucrose solution(s)/cushion components and keep them chilled on ice until use.
c. Add certain supplements (for example, inhibitors/reducing agents) immediately before use when indicated.
d. If magnetic cleanup is used, prepare the bead-based nuclei enrichment reagents according to the manufacturer’s instructions and maintain them at recommended temperatures.
3. **Pre-stage tubes and plan workflow timing**

a. Label all tubes for homogenization, gradient loading, nuclei recovery, wash steps, and final resuspension before beginning.
b. Plan the processing sequence so homogenization, gradient setup, and centrifugation can be completed without interruption.
c. Plan the workflow so that, once homogenization begins, samples proceed to density-based cleanup without unnecessary delays. Avoid holding homogenized samples on ice for extended periods before cleanup, as this can increase debris accumulation and myelin carryover.
4. **Prepare for nuclei counting and quality assessment**

a. Set up materials for nuclei counting and microscopic inspection (for example if used a counting chamber or a stain).
b. Define parameters for acceptable nuclei integrity and debris burden before processing experimental samples.
c. Confirm nuclei concentration and loading requirements for the intended downstream snRNA-seq platform.
5. **Prepare tissue samples**

a. Dissect the adult mouse hippocampus or cerebellum from the brain and briefly rinse the tissue in ice-cold PBS to remove excess blood.
b. Transfer the rinsed tissue immediately into ice-cold NIB (Nuclei Isolation Buffer) for downstream processing.
c. Record tissue mass before homogenization to guide input-specific handling and calculations for expected recovery. CRITICAL: Dissect tissue only after all reagents and equipment are prepared. After the brief PBS rinse, transfer tissue promptly into ice-cold NIB and keep it cold to minimize sample degradation.

### Innovation

This protocol presents a benchtop-compatible nuclei isolation workflow optimized for myelin-rich adult mouse brain tissue (hippocampus and cerebellum) for downstream snRNA-seq. Standard brain nuclei workflows typically combine mechanical dissociation with density-based cleanup to reduce myelin and debris, and many sucrose-gradient implementations rely on ultracentrifugation and careful gradient handling (1–5). These features can increase hands-on complexity and may limit use in laboratories without ultracentrifugation equipment (3,5). Our workflow incorporates three practical modifications. First, tissue can be homogenized using a tube-and-pestle format rather than a conventional Dounce setup, providing a less labor-intensive option for tissue disruption (1–3). Second, nuclei are purified using a low-volume (2 mL) sucrose gradient compatible with a standard benchtop centrifuge, enabling nuclei recovery by pelleting rather than interphase collection (3,7). Third, an optional magnetic bead–based nuclei enrichment step can be added after pelleting to further reduce visible myelin/debris carryover during cleanup.

Under the tested conditions, this workflow supports the generation of nuclei suspensions suitable for downstream 10x Genomics Flex and PARSE WT library preparation from adult mouse hippocampus and cerebellum. Further optimization may be required for other brain regions or species.

### Protocol Preparation

**Timing:** 20–30 min

1. **Pre-chill workspace and instruments**

1.1 Set centrifuges to 4°C and pre-cool the swinging-bucket rotor.
1.2 Place a tube rack and (if dissecting fresh tissue) a pre-chilled aluminum block on ice.
2. **Stage consumables on ice (RNase-minimizing setup)**

2.1 Pre-chill low-binding tubes (1 or 2 mL) and set out low-retention tips (use wide-bore/trim tips when handling nuclei to reduce shear).
2.2 Place disposable pestles (and/or a 2 mL Dounce) on ice.
2.3 Place an unopened sterile 30 µm filter/mesh on ice and open it immediately before filtration.
3. **Prepare ice-cold buffers (fresh additives)**

3.1 Prepare/aliquot NIB, Nuclei Resuspension Buffer (NSB), Adjusted Sucrose Solution, and Magnetic Resuspension Buffer (MRB) and keep on ice.
3.2 Add DTT and protease inhibitor to the NIB immediately before use.
3.3 Add RNase inhibitor immediately before use to buffers where indicated (e.g., NSB/MRB; optional in NIB).
4. **Set up sucrose cushion workflow**

4.1 Pre-aliquot 500 µL Adjusted Sucrose Solution per sample (for the cushion).
4.2 Plan the sample mix: you will later add 900 µL Adjusted Sucrose Solution to the nuclei suspension prior to layering.
5. Prepare magnetic cleanup station (if using Anti-Nucleus MicroBeads)

5.1 Place the MiniMACS™ Separator on the MACS MultiStand, insert an MS column, and pre-chill the required buffer(s).
5.2 Stage a timer and collection tubes for the labeling/incubation and washes.

**CRITICAL:** Keep tissue, buffers, and tools on ice throughout to preserve nuclear integrity and minimize RNA degradation.

## KEY RESOURCES TABLE

| REAGENT or RESOURCE | SOURCE | IDENTIFIER |
| --- | --- | --- |
| Sucrose | Research Products International | S24060-1000.0 |
| Potassium chloride (KCl) | Research Products International | P41025-500 |
| Magnesium chloride hexahydrate (MgCl<sub>2</sub>·6H<sub>2</sub>O) | Fisher Chemical | M33-500 |
| Tris-HCl | Research Products International | T60050-500.0 |
| 0.5 M EDTA, pH 8.0 | Invitrogen (Thermo Fisher Scientific) | AM92660G |
| Phosphate-buffered saline (PBS), pH 7.4 | Gibco | 10010-023 |
| IGEPAL CA-630 | Sigma-Aldrich | 18896-50ML |
| Dithiothreitol (DTT) | Research Products International | D11000-5.0 |
| Protease inhibitor cocktail, EDTA-free | Thermo Scientific | 78425 |
| RNase inhibitor | Thermo Scientific | EO0381 |
| Bovine serum albumin (BSA), 10% in DPBS | Sigma-Aldrich | A1595-50ML |
| Nuclease-free water | Research Products International | 248700-500.0 |
| Anti-Nucleus MicroBeads | Miltenyi Biotec | 130-132-997 |
| MS columns | Miltenyi Biotec | 130-042-201 |
| Acridine Orange (AO) | Invitrogen (Thermo Fisher Scientific) | A-3568 |
| Propidium iodide (PI) | Fisher Scientific | P3566 |

## MATERIALS AND EQUIPMENT

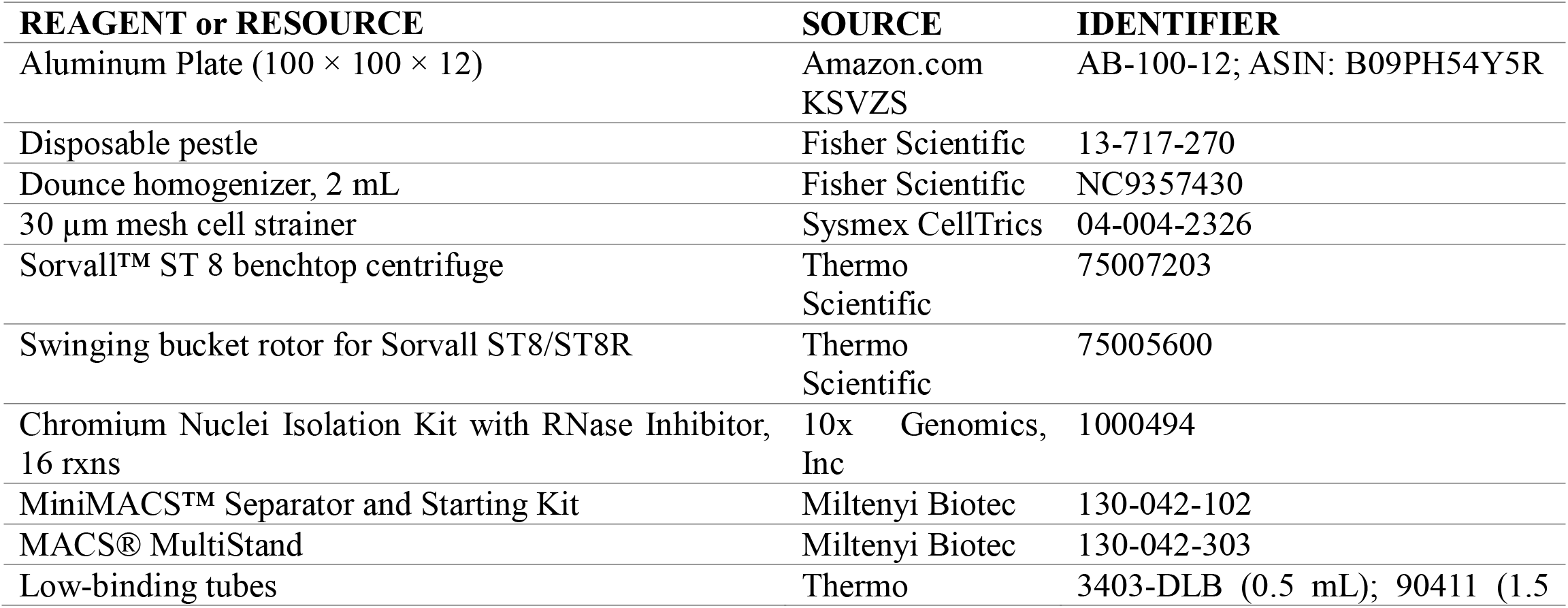
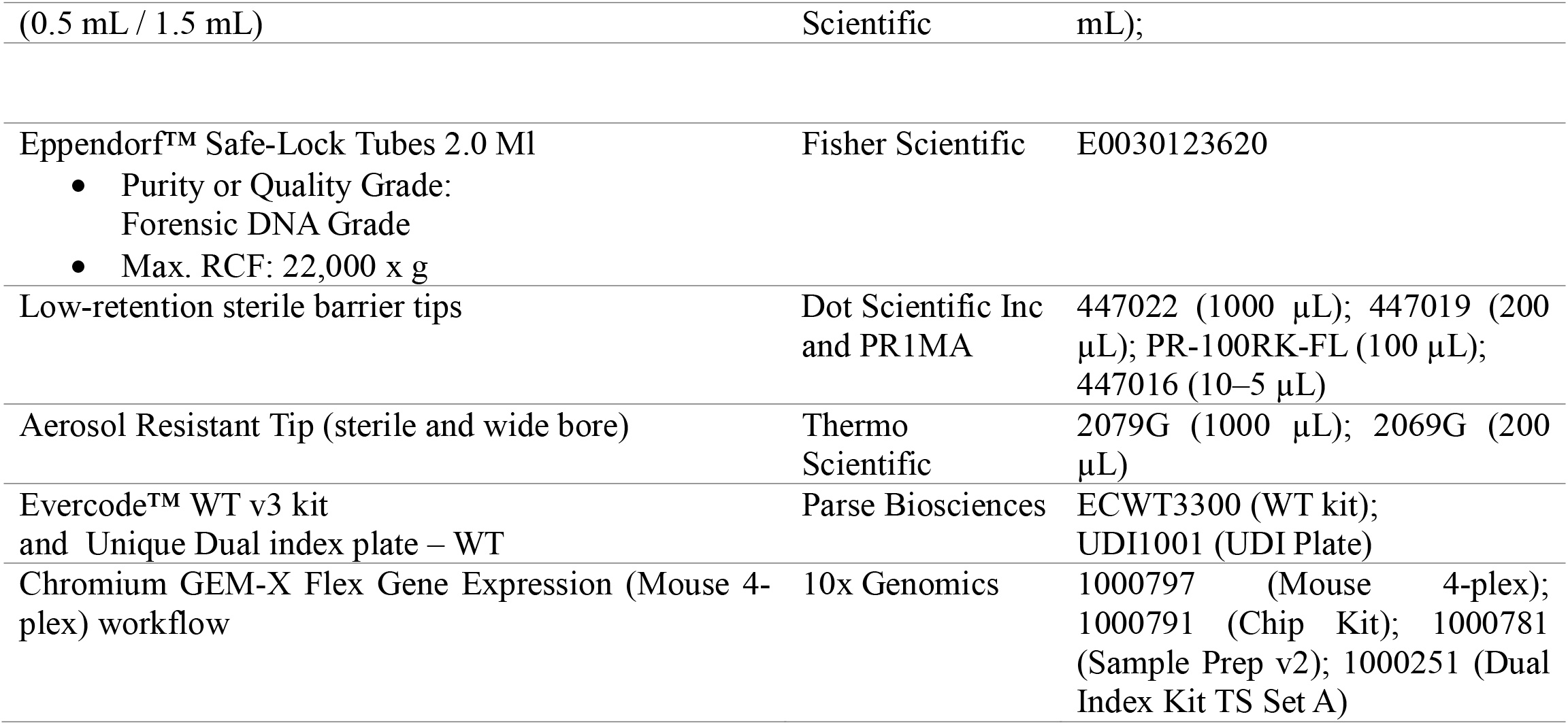

## BUFFERS

Keep completed working buffers on ice during the procedure. Prepare stock solutions and store them at the temperatures indicated below. Keep temperature-sensitive additives (including DTT, protease inhibitor, and RNase inhibitor) chilled during buffer preparation and add them immediately before use.

### Nuclei Isolation Buffer (NIB)

| Component | Stock concentration | Final concentration | Notes |
| --- | --- | --- | --- |
| Sucrose | 2.5 M | 250 mM | Prepare in nuclease-free water; store at 4°C |
| KCl | 1 M | 25 mM | Prepare in nuclease-free water; store at 4°C |
| MgCl<sub>2</sub> | 1 M | 5 mM | Prepare in nuclease-free water; store at 4°C |
| Tris-HCl, pH 7.4 | 1 M | 10 mM | Prepare in nuclease-free water; store at 4°C |
| DTT | 1 M | 1 mM | Add immediately before use, freshly prepare right before protocol start; keep chilled during preparation |
| Protease inhibitor (EDTA-free) | 100× | 1× | Add immediately before use; keep chilled during preparation. |
| RNase inhibitor | 0.2 U/µL | 0.2 U/µL | Add immediately before use; keep chilled during preparation. |
| Nuclease-free water | — | to volume | Mix gently |
| IGEPAL | 10% | 0.1–0.3% | Add last; mix gently |

### Nuclei Resuspension Buffer (NSB)

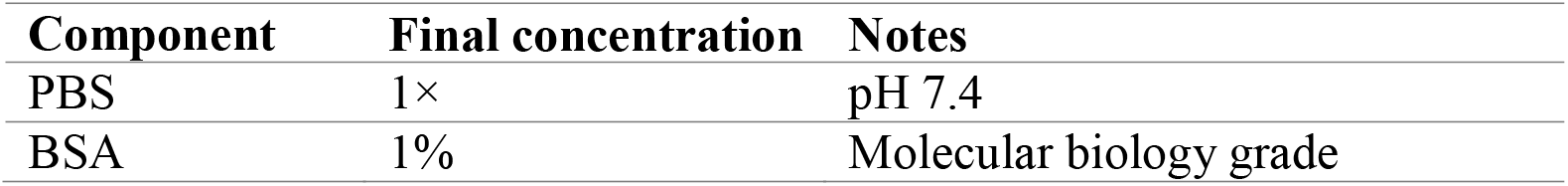
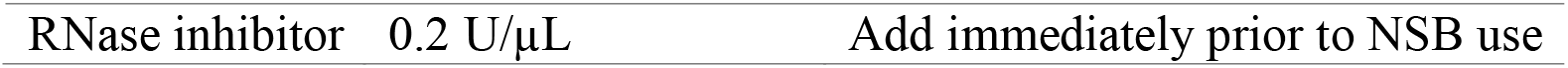

### Adjusted Sucrose Solution (Adj.S)

| Component | Stock concentration | Working concentration | Notes |
| --- | --- | --- | --- |
| Sucrose | 2.5 M | 1.8 M | Prepare in nuclease-free water; store at 4°C. A 2.0 M condition was also used successfully in selected experiments; exact concentration is specified where applicable. |
| MgCl<sub>2</sub> | 1 M | 3 mM | Prepare in nuclease-free water; store at 4°C |
| Tris-HCl, pH 8.0 | 1 M | 10 mM | Prepare in nuclease-free water; store at 4°C |
| Nuclease-free water | — | to volume |  |

### Magnetic Resuspension Buffer (MRB)

| Component | Final composition | Notes |
| --- | --- | --- |
| PBS, pH 7.2 | 1× | Store at 4°C |
| BSA | 1% | Keep on ice |
| EDTA, pH 8.0 | 0.5 mM |  |
| RNase inhibitor | 0.2 U/μL | |

### AO/PI staining solution

Aliquots of the completed 2× AO/PI staining solution may be stored at **2–8°C** protected from light. Do not freeze, as freezing can alter the solution.

| Component | Final composition | Notes |
| --- | --- | --- |
| PBS | 1× | Use sterile, filtered PBS. |
| Acridine Orange (AO) | 10 μg/mL (in 2×) | Prepare a 1 mg/mL AO intermediate by mixing 10 μL of 10 mg/mL AO stock with 90 μL PBS. |
| Propidium iodide (PI) | 200 μg/mL (in 2×) | Use 1 mg/mL PI stock. |
| PBS (to final volume) | — | Add PBS to bring the final 2× staining solution volume to 100 μL. |

### To prepare 100 µL of 2× AO/PI staining solution

Combine 78 µL PBS, 2 µL of the 1 mg/mL AO intermediate, and 20 µL of 1 mg/mL PI stock. Mix gently and protect from light.

### To stain nuclei

Mix the completed 2× AO/PI staining solution 1:1 with sample (for example, 5 µL sample + 5 µL 2× stain), incubate for 1–5 min in the dark at room temperature, then load the stained sample into the counting chamber or instrument for analysis.

## STEP-BY-STEP METHOD DETAILS

### Nuclei isolation and purification from mouse hippocampus

**Timing:** ∼2.5–3.5 h (if magnetic enrichment is omitted for unusually clean preparations); ∼3–4.5 h (with magnetic enrichment, recommended

**CRITICAL:** Keep all buffers, tubes, and tissue ice-cold throughout. Work quickly to minimize RNA degradation, especially before nuclei are isolated and transferred into RNase inhibitor–containing buffer. RNA degradation risk is reduced after this step, but samples should still be kept cold and processed promptly.

#### 1. Tissue collection and mincing

**Timing:** 10–15 min

1. Dissect hippocampi rapidly (∼3–4 min) on a pre-chilled aluminum block.
2. For nuclei isolation, transfer tissue directly into ice-cold Nuclei Isolation Buffer (NIB). For long-term storage, snap-freeze tissue in liquid nitrogen and store in -80 °C.
3. Keep tissue fully submerged in NIB. Using pre-chilled tools, mince into ∼1 mm³ pieces. As a guide, use 1:10 (w/v) tissue:buffer (e.g., 100 mg tissue in 1 mL NIB) and adjust as needed to keep tissue fully submerged.

**NOTE:** If working from frozen tissue, minimize thaw time by transferring directly into cold NIB.

#### 2. Mechanical homogenization

Choose one of the following homogenization options. For both homogenization options, use approximately 1:10 (w/v) tissue:NIB as the default starting ratio. Adjust the absolute buffer volume only as needed to keep tissue fully submerged and to allow smooth mechanical dissociation without excessive crowding or dilution.

### Option A: Tube and pestle homogenization

**Timing:** 5–10 min

1. Transfer minced tissue to a pre-chilled 2 mL low-binding tube containing ice-cold NIB (use enough to fully submerge tissue (typically ∼1 mL total volume in a 2 mL tube for ∼50–100 mg tissue) maintaining ∼1:10 w/v tissue:buffer as a guide).
2. Homogenize with a sterile disposable pestle using 10–15 slow, gentle strokes (∼1–2 seconds per stroke from bottom to top of the tube). Place the tube back on ice between every few strokes to re-chill the tube and keep on ice immediately after homogenization. Avoid aggressive strokes or crushing tissue against the tube wall/bottom, as this can rupture nuclei and increase foaming in the tube.

**NOTE:** Do not vortex or crush tissue against the tube wall/bottom. If the slurry does not pass smoothly through a 1 mL tip, use wide-bore or trimmed tips and continue gentle strokes until no large fragments remain.

### Option B: Dounce homogenization

**Timing:** 10–15 min

1. Transfer minced tissue into a pre-chilled 2 mL Dounce homogenizer containing ice-cold NIB, using approximately 1:10 (w/v) tissue:NIB as the starting ratio. In practice, the total volume may be increased as needed for the Dounce format to keep tissue fully immersed and allow smooth pestle movement.
2. Homogenize on ice with the loose pestle (A) using 5–10 slow, gentle strokes (∼1–2 s per stroke, from top to bottom and back to the start position). Place the homogenizer on ice between every few strokes to re-chill.
3. Continue with the tight pestle (B) using ∼10 slow, controlled strokes (∼1–2 s per stroke). Place the homogenizer on ice between every few strokes to re-chill and keep on ice immediately after homogenization.

**NOTE:** Do not force tissue through resistance or use aggressive plunging/grinding

**CRITICAL:** For either homogenization method, use slow, controlled strokes, keep samples and ice-cold tools, and place the tube or homogenizer back on ice between every few strokes to minimize shear, preserve nuclear integrity, and reduce foaming. Avoid aggressive homogenization.

#### 3. Clarification, filtration, and resuspension

**Timing:** 20–25 min

1. Transfer homogenate to a pre-chilled 2 mL low-binding tube and centrifuge at ∼500 × g for 5 min at 4°C.
2. Carefully discard the supernatant without disturbing the pellet
3. Gently resuspend the pellet in 500 µL ice-cold NSB using wide-bore tips and keep on ice.
4. Filter the resuspended pellet through a 30 µm mesh into a pre-chilled low-binding tube.
5. Rinse the filter with an additional 500 µL ice-cold NSB, collect the rinse in the same tube as the filtrate, and keep on ice.
6. Centrifuge the combined filtrate at ∼500 × g for 5 min at 4°C.
7. [Before resuspending, read critical point below] Carefully discard the supernatant and gently resuspend the pellet in 500 µL ice-cold NSB. Keep on ice until proceeding to the next step.

**CRITICAL:** The resuspension volume should be adjusted based on the residual buffer already present with the pellet. Small pellets (for example, ∼20 mg hippocampus) may retain only ∼20–30 µL residual volume, so add ∼470–480 µL NSB to reach a final volume of 500 µL. Larger pellets (for example, cerebellum) may retain substantially more residual volume, so correspondingly less NSB may be needed (for example, ∼300–400 µL added). In all cases, resuspend to a final total volume of 500 µL.

#### 4. Sucrose cushion and layering of sample

**Timing:** 10–15 min

1. With your pipette tilted ∼10-15**°** to minimize layer disruption, add 500 µL Adjusted Sucrose Solution (Adj.S) to the bottom of a sterile 2 mL low-binding Eppendorf tube to form the cushion (Fig. 1a). We recommend 1.8 M as the default starting condition; 2.0 M was also used successfully under the tested conditions.
2. To the nuclei suspension (from step 3.7, ∼500 µL), add 900 µL Adj.S (Fig. 1a). Slowly pipette the Adj.S to a final volume of 1.4 mL and incubate on ice for 1–2 min.

**Figure 1.**
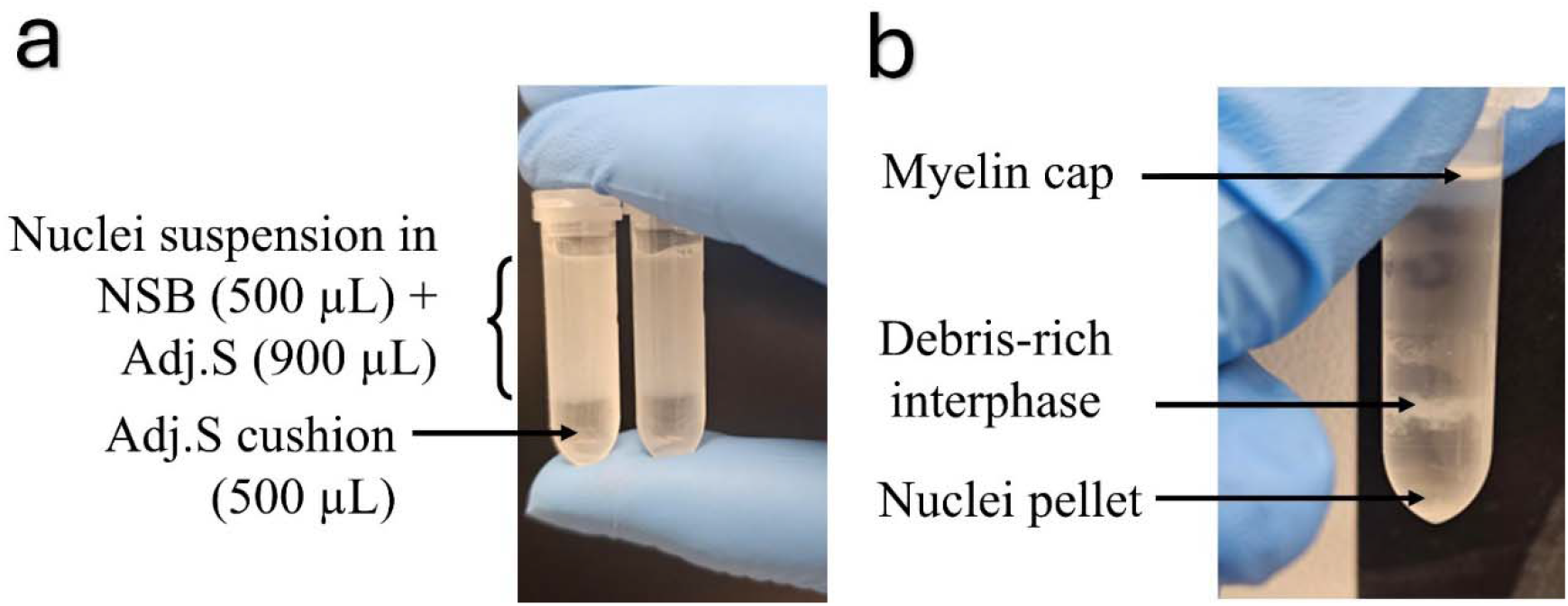
Sucrose cushion setup and phase separation during nuclei purification from adult mouse brain. **(a) Cerebellum (sucrose cushion setup).** Representative setup for density pelleting: resuspended nuclei in nuclei suspension buffer (NSB; 500 µL) were mixed with adjusted sucrose solution (Adj.S; 900 µL) and carefully layered onto a 500 µL Adj.S cushion at the bottom of a 2 mL tube. The increased turbidity is indicative of myelin-rich tissue. **(b) Hippocampus (post-sucrose separation).** Representative tube after density centrifugation showing the typical phase structure, including a myelin cap (top), debris-rich interphase (middle band), and nuclei pellet (bottom; often faint). For optimal recovery, remove the myelin cap first, aspirate the interphase without disturbing the pellet, and leave a small residual volume above the pellet to minimize nuclei loss. ***Note:*** *Panels (a) and (b) show representative preparations from different tissues (cerebellum vs hippocampus) and are not the same sample*.

Carefully layer the 1.4 mL mixture on top of the sucrose cushion by tilting the tube 10–15**°** and dispensing the mixture along the tube wall, then return the tube upright onto ice.

**CRITICAL:** Layers are delicate, do not disturb the interface between layers.

#### 5. Density centrifugation and myelin removal

**Timing:** 55–60 min

1. Centrifuge at 13,000 × g for 45 min at 4°C.
2. After centrifugation (Fig. 1b), you should see;
3. a white myelin cap at the top,
4. a clearer phase below,
5. nuclei pelleted near the bottom (pellet may be small/not visible).
6. Remove the myelin cap first by gently sliding it along the tube wall and lifting/aspirating the cap (detailed in troubleshoot Problem 7). **NOTE:** Myelin can clog tips—sliding is often easier than aspiration.
7. Carefully aspirate the remaining supernatant (including the interphase) without disturbing the pellet. Because the nuclei pellet may be faint or not visible, aspirate slowly and leave ∼100–200 µL above the pellet to minimize nuclei loss.

**OPTIONAL:** For cerebellar samples, users may collect the bottom pellet only to prioritize purity over yield. This can reduce debris/myelin carryover and may lessen the need for downstream magnetic enrichment but typically lowers total nuclei recovery.

#### 6. Wash nuclei and prepare final suspension

**Timing:** 20–30 min

1. Add 500 µL ice-cold NSB to the pellet and gently resuspend. Transfer to a fresh pre-chilled 2 mL low-binding Eppendorf tube.
2. Centrifuge at ∼500 × g for 5 min at 4°C. Discard the supernatant.
3. Resuspend nuclei pellet in 500 µL NSB.
4. Filter through a 30 µm mesh, rinse with an additional 500 µL NSB. To ensure residual sucrose is removed for optimal downstream library preparation, repeat steps 1 – 3 two to three times.
5. Centrifuge at ∼500 × g for 5 min at 4°C. Discard the supernatant.
6. Resuspend the final nuclei pellet in 70–100 µL MRB. Count nuclei at this stage before proceeding. Proceed to downstream snRNA-seq only if the preparation is nuclei-dominant by microscopy, with predominantly round nuclei-sized particles, minimal large debris, and little obvious clumping, and if the final nuclei concentration is sufficient for the intended downstream platform. AO/PI-based counting should be interpreted as a supportive QC readout rather than a strict viability metric for isolated nuclei.

**CRITICAL:** Keep nuclei on ice throughout these wash steps. Nuclei can be held on ice for ∼30–60 min before downstream processing but proceed as soon as possible for best RNA quality. Prior nuclei counting is required before magnetic separation.

### Magnetic enrichment using Anti-Nucleus MicroBeads

Perform this cleanup step to obtain the final sequencing-ready nuclei preparation, particularly for myelin-rich samples. In unusually clean preparations after sucrose pelleting, this step may be omitted, but downstream sample quality should be confirmed by microscopy and counting.

#### 7. Magnetic labeling

**Timing:** 45-50 min

1. Keep nuclei in ice-cold MRB. Recommended input is ≤1 × 10□nuclei per MS column.
2. Transfer up to 1 × 10□nuclei into a 1.5 mL low-binding Eppendorf tube and centrifuge at 500 × g for 5 min at 4°C. Discard the supernatant.
3. Resuspend pellet in 450 µL MRB
4. Add 50 µL Anti-Nucleus MicroBeads per 1 × 10□nuclei.
5. Mix gently and incubate at 4°C for 15–30 min with gentle shaking or rocking.

**CRITICAL:** Mix gently by inverting the tube—do not vortex.

#### 8. Wash labeled nuclei

**Timing:** 10–15 min
1. Add 1–2 mL MRB (This is a wash step to remove unbound/excess MicroBeads (and reduce background/non-specific carryover) before loading onto the MS column)
2. Centrifuge at 500 × g for 5 min at 4°C. Discard the supernatant.
3. Resuspend nuclei in 500 µL MRB per MS column.

## 9. Magnetic separation (MS columns + MiniMACS)

**Timing:** 15–25 min

1. Place MS column in the MiniMACS separator and pre-rinse with 500 µL MRB.
2. Apply labeled nuclei to the column. Collect flowthrough (negative fraction; unlabeled/non-retained nuclei) if performing optional QC or troubleshooting; otherwise discard.
3. Wash column 3× with 500 µL MRB each time, letting it fully run through every time.
4. Remove column from magnet and place over a new 2 mL low-binding Eppendorf tube.
5. Elute enriched nuclei by adding 1 mL MRB and pushing through with the plunger (provided with the kit).
6. Centrifuge eluate at 500 × g for 5 min at 4°C. Discard the supernatant.
7. Resuspend nuclei pellet in 50–100 µL NSB (adjust based on expected yield). Keep on ice.

## EXPECTED OUTCOMES

### Optimization of nuclei purification in myelin-rich adult mouse brain

This protocol is optimized to generate intact, debris-reduced nuclei suspensions from myelin-rich adult mouse brain for downstream single-nucleus RNA-seq (snRNA-seq). Successful preparations are characterized by (i) a nuclei-dominant particle population by microscopy, (ii) low carryover of myelin/debris after density-based cleanup, and (iii) compatibility with standard snRNA-seq library construction and quality-control (QC) filtering.

We first tested an OptiPrep density gradient (29–50%) to remove myelin/debris from adult mouse cerebellum nuclei preparations (∼20 mg input). This condition had very low recovery of usable nuclei and had poor nuclei integrity as evidenced by dye-based counting: total nuclei were 4.56 ×□10□± 4.80 × 10□nuclei/mL, and all detected events were dye-positive/compromised (1.01 × 10□nuclei/mL; 0.0% stain-negative; 45 total nuclei counted, 0 stain-negative/45 stain-positive), with a mean reported particle diameter of 8.3 µm (Fig. 2a). Microscopy corroborated these measurements, showing sparse particles dominated by dye-positive nuclei and substantial background debris, consistent with extensive damage and/or non-nuclear contamination (Fig. 2a). Accordingly, AO/PI-based automated counting was used here as a practical QC readout for nuclei concentration and sample composition, and its interpretation in isolated nuclei should be considered an adaptation for nuclei preparations rather than a validated live/dead viability standard. Because plasma membranes are intentionally disrupted during nuclei isolation, liberated nuclei typically appear PI-dominant (red), whereas residual intact or incompletely lysed whole cells typically appear AO-dominant (green). We therefore use this color pattern as a qualitative indicator of nuclei liberation versus intact-cell carryover and assess sample quality primarily by microscopy (nuclear morphology/debris) and snRNA-seq QC metrics, rather than by automated AO/PI “viability” percentages.

**Figure 2.**
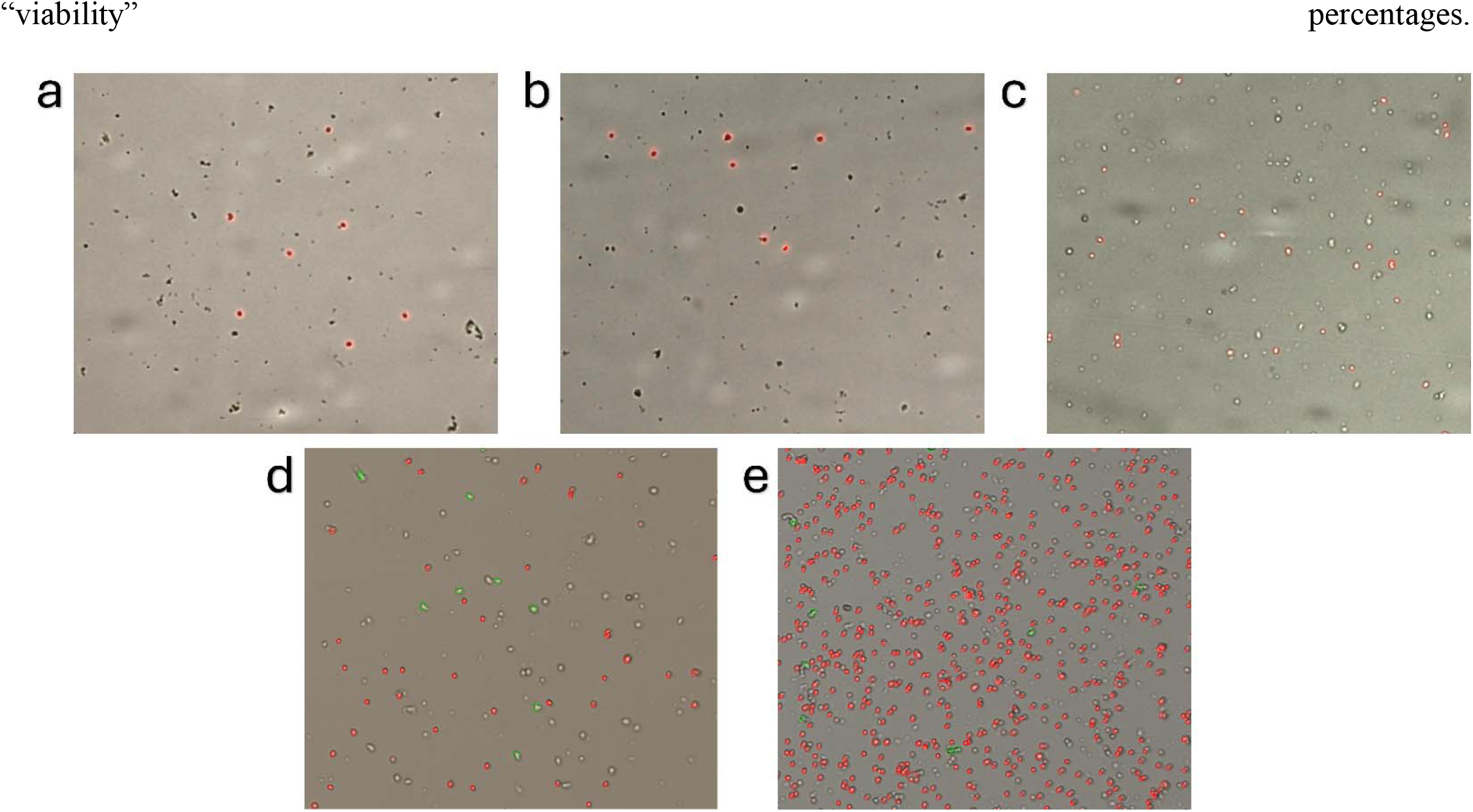
Representative fluorescence images from dye-based nuclei assays during exploratory optimization of nuclei cleanup from adult mouse brain tissue. (a) Cerebellum (30 mg), OptiPrep 29–50%: 4.56 × 10□± 4.80 × 10□nuclei/mL; 0/45 stain-negative events detected. (b) Cerebellum (35 mg), OptiPrep 29%: 1.37 × 10□± 5.31 × 10□nuclei/mL; 6/81 stain-negative events detected. (c) Hippocampus (19 mg), Chromium Nuclei Isolation Kit (10x Genomics): 2.03 × 10□± 1.60 × 10□nuclei/mL; (d) Cerebellum (22 mg), sucrose pelleting (4–20 µm gate): 2.68 × 10□± 5.93 × 10³, Adj.S = 1.8 M (e) Cerebellum (70 mg), sucrose pelleting (4–20 µm gate): 1.04 × 10□± 2.23 × 10□nuclei/mL, Adj.S = 1.8 M Cell/nuclei concentrations are reported as mean ± SD cells/mL of original sample after correction for the 1:2 AO/PI dilution. Colored overlays indicate instrument-classified isolated nuclei (red; PI-dominant) and residual intact/unlysed cells (green; AO). Because tissue region, input mass, and workflow were not matched across all panels, these images are presented as representative optimization examples rather than a controlled head-to-head comparison.

**Figure 3.**
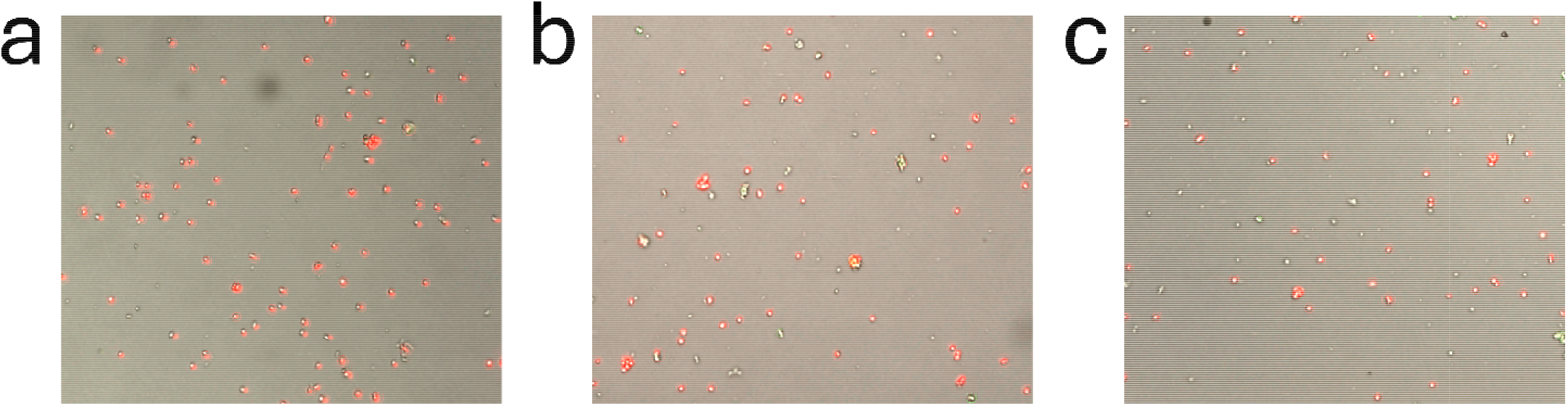
Representative fluorescence images after magnetic nuclei enrichment of adult mouse brain nuclei preparations. (a) Cerebellum, sucrose pelleting followed by magnetic nuclei enrichment: before enrichment, 5.50 × 10^7^ ± 6.10 × 10^6^ nuclei/mL; after enrichment, 2.13 × 10^6^ ± 6.07 × 10^5^ nuclei/mL, corresponding to ∼3.9% recovery of pre-enrichment nuclei. (b) Hippocampus, sucrose pelleting followed by magnetic nuclei enrichment: before enrichment, 2.26 × 10^6^ ± 1.77 × 10^6^ nuclei/mL; after enrichment, 1.72 × 10^6^ ± 5.44 × 10^5^ nuclei/mL, corresponding to ∼76% recovery of pre-enrichment nuclei. (c) Hippocampus, 10x Chromium Nuclei Isolation Kit followed by magnetic nuclei enrichment: before enrichment, 5.89 × 10^6^ ± 2.46 × 10^6^ nuclei/mL; after enrichment, 1.18 × 10^6^ ± 3.05 × 10^5^ nuclei/mL, corresponding to ∼20% recovery of pre-enrichment nuclei. Nuclei concentrations are reported as mean ± SD nuclei/mL of original sample after correction for the 1:2 AO/PI dilution. Colored overlays indicate instrument-classified isolated nuclei and residual intact/unlysed cells. These images illustrate that magnetic enrichment reduced visible debris and improved sample cleanliness, although recovery varied substantially by tissue region and upstream isolation workflow.

We next evaluated a lower-density OptiPrep condition (29% only) using cerebellum at a larger input mass (35 mg; Fig. 2b). Under these conditions, total nuclei concentration increased to 1.37 × 10□± 5.31 × 10□ nuclei/mL, including 1.34 × 10^4^ stain-negative nuclei/mL and 1.68 × 10^5^ stain-positive nuclei/mL, with an average reported particle size of 7.8 µm. Microscopy again showed few nuclei-like particles and substantial debris (Fig. 2b). We then applied the Chromium Nuclei Isolation Kit workflow to adult hippocampus (19 mg input; Fig. 2c). This preparation yielded 2.03 × 10□± 1.60 × 10□nuclei/mL, with all detected events classified as stain-positive/compromised and a small average reported particle size (3.8 µm), together with abundant small particles and background debris by microscopy.

Because residual lipid/myelin burden remained prominent in these early optimization conditions, we next implemented a benchtop sucrose-gradient pelleting workflow in cerebellum and quantified recovery using a 4– 20 µm size gate. With a 22 mg cerebellum sample, sucrose pelleting yielded 1.69 × 10□± 2.24 × 10□ nuclei/mL (Fig. 2d). When applied to a larger cerebellum input (70 mg), the same workflow yielded 1.04 × 10□ ± 2.23 × 10□nuclei/mL (Fig. 2e). Microscopy in both preparations showed dense fields of nuclei-sized particles with reduced large debris. Because the data shown in Figure 2 were derived from varied brain regions, tissue masses, and workflows, these results should be interpreted as representative optimization examples rather than a controlled comparison of method performance. Because dye-based classification relies on algorithmic assignment (signal plus size/shape heuristics), “intact cell” calls may reflect residual small cells and/or size-based misclassification rather than true intact cells. Consistent with minimal cellular carryover, microscopy showed a dense field of nuclei-sized particles with minimal large debris (Fig. 2e), supporting sucrose pelleting’s robust recovery from higher mass, higher myelin-rich inputs. Taken together, these optimization experiments informed the final workflow and parameter set used in this protocol. Because OptiPrep-based conditions and the Chromium Nuclei Isolation Kit used as a standalone isolation method yielded low-recovery, debris-rich preparations under our tested conditions, we advanced the benchtop sucrose-pelleting workflow for further development. The reported results presented here are based on the sucrose-pelleting conditions that performed successfully in these optimization experiments, including detergent-assisted homogenization with 0.2% IGEPAL, low-volume sucrose pelleting in 2 mL tubes, and centrifugation at 13,000 × g for 45 min at 4°C. Because residual debris/myelin carryover could persist after pelleting, particularly in cerebellar preparations, magnetic enrichment was incorporated as the recommended downstream cleanup step for sequencing-ready nuclei suspensions. Although 0.2% IGEPAL worked for the tissues evaluated here, detergent requirements may vary across tissue types and sample states and may require further optimization in other contexts.

### Magnetic separation improves debris cleanup

Magnetic separation improves debris cleanup with distinct consequences for apparent nuclei yield across workflows. We evaluated the effect of magnetic enrichment on nuclei yield and composition using both sucrose pelleting–based and Chromium Nuclei Isolation Kit–based workflows, quantifying nuclei from a standardized 100 µL resuspension volume using an automated counter (DeNovix). For sucrose pelleting–based cerebellar preparations, samples yielded 5.50 × 10□± 6.10 × 10□nuclei/mL after sucrose pelleting but only 2.13 × 10□± 6.07 × 10□nuclei/mL after magnetic enrichment, corresponding to substantially lower recovery (∼3.9% of pre-enrichment nuclei) and improved debris removal. For sucrose pelleting–based hippocampal preparations, samples yielded 2.26 × 10□± 1.77 × 10□nuclei/mL prior to cleanup and 1.72 × 10□ ± 5.44 × 10□nuclei/mL following magnetic enrichment (∼76% recovery). In this case, sucrose pelleting alone already produced largely myelin- and debris-free suspensions, and magnetic enrichment was used mainly to quantify potential losses rather than to correct major contamination (Supplementary Fig. 1a).

In contrast, for the Chromium Nuclei Isolation Kit–based hippocampal workflow, pre-enrichment suspensions contained abundant debris and small particles that inflated automated “nuclei” counts. Apparent pre-enrichment counts were 5.89 × 10□± 2.46 × 10□nuclei/mL, but this value overestimates true nuclei abundance because non-nuclear material was frequently classified as nuclei (Supplementary Fig. 1b). After magnetic cleanup, samples yielded 1.18 × 10□± 3.05 × 10□nuclei/mL, corresponding to ∼20% of the pre-enrichment count but a much cleaner preparation. Thus, for Chromium Nuclei Isolation Kit–based hippocampal samples, the low apparent “recovery” primarily reflects removal of debris rather than loss of intact nuclei, and pre-enrichment counts should not be directly compared to those from sucrose-pelleted hippocampus without considering this difference in debris burden.

Importantly, the elevated pre-cleanup yield observed in cerebellar preparations likely reflects substantial debris carryover during sucrose pelleting. Adult cerebellar tissue contains extremely dense populations of small granule neurons, generating abundant fine particulate material during homogenization (8,9). Under sucrose-based cleanup conditions, myelin is enriched toward the upper layer, whereas nuclei and fine debris may remain distributed between the pellet and the debris-rich intermediate fraction (1,3,7). Accordingly, recovery strategies that include selected intermediate material can increase apparent yield but also increase non-nuclear carryover.

In sucrose-based nuclei isolation workflows, myelin and debris are typically separated from nuclei by density behavior during centrifugation, but clean partitioning can remain challenging in lipid-rich brain tissue and debris-rich preparations (3). Accordingly, to maximize recovery from cerebellar preparations, we collected the bottom pellet together with selected material from the debris-enriched intermediate fraction; this increased apparent pre-cleanup yield but also produced debris-rich suspensions that required additional downstream cleanup (Fig. 4). Consistent with this interpretation, magnetic enrichment reduced non-nuclear carryover and yielded a cleaner final nuclei population, albeit with reduced overall recovery in cerebellar samples. This pattern is consistent with the purity-yield tradeoff commonly encountered in nuclei isolation from myelin-rich brain tissue, where more stringent cleanup improves sample purity at the cost of nuclei recovery (3). Together, these results indicate that magnetic enrichment improves nuclei purity across workflows, but recovery remains strongly influenced by tissue-specific properties, particularly in cerebellum.

**Figure 4.**
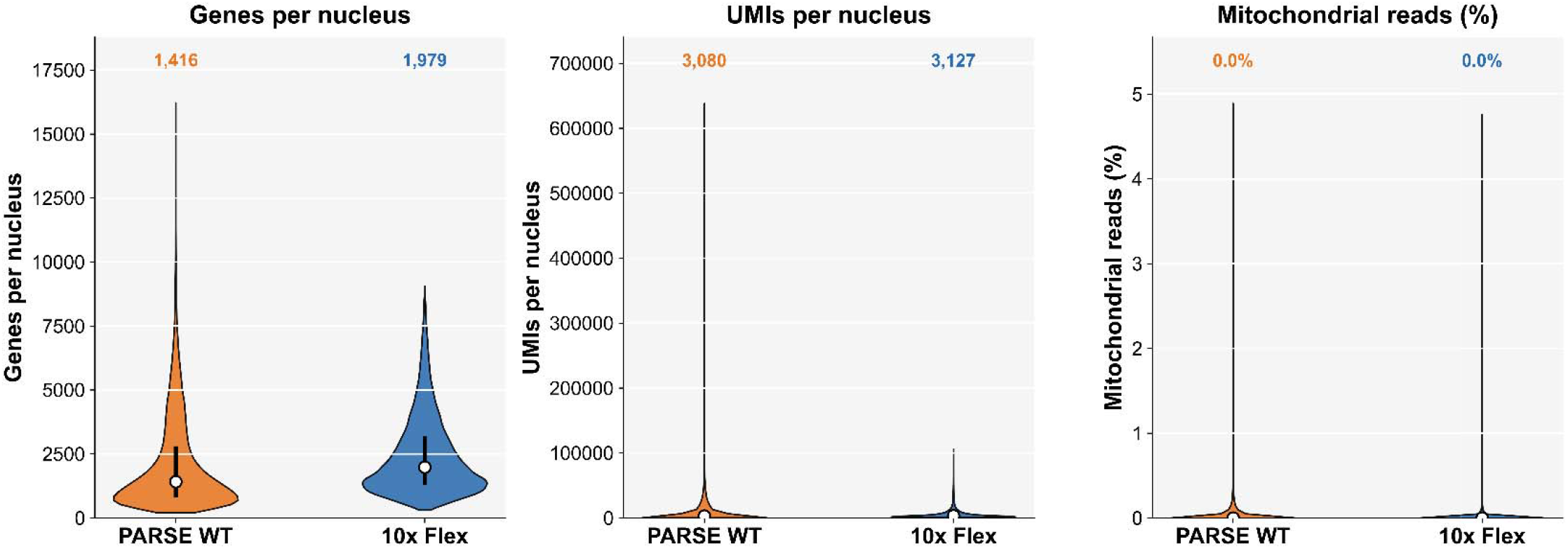
Aggregate post-QC overview of snRNA-seq quality metrics across PARSE WT and 10x Flex datasets. Violin plots show QC-retained nuclei aggregated across PARSE WT hippocampal samples prepared using the Chromium Nuclei Isolation Kit followed by magnetic enrichment (n = 20 samples) and 10x Flex cerebellar samples prepared using sucrose pelleting followed by magnetic enrichment (n = 12 samples). Left, genes detected per nucleus (n_genes_by_counts); middle, UMIs per nucleus (total_counts); right, mitochondrial read fraction (pct_counts_mt). Numbers above violin plots indicate median values for each dataset. Because tissue type, isolation workflow, and sequencing chemistry differed between datasets, this figure is intended as a feasibility/QC overview rather than a controlled head-to-head comparison of platform performance.

### Downstream snRNA-seq results

To evaluate whether nuclei generated by the tested workflows supported downstream snRNA-seq analysis, we examined post-QC distributions of core per-nucleus metrics, including genes detected per nucleus (n_genes_by_counts), UMIs per nucleus (total_counts), and mitochondrial read fraction (pct_counts_mt) (Fig. 4). Libraries were generated using either the Evercode™ WT v3 single-cell/single-nucleus RNA-seq workflow (Parse Biosciences; ECWT3300; referred to here as PARSE WT), a split-pool combinatorial barcoding–based platform, or the Chromium GEM-X Flex Gene Expression workflow (10x Genomics; referred to here as 10x Flex), a probe-based fixed-RNA whole-transcriptome workflow (10–12). The PARSE WT dataset comprised hippocampal samples prepared using the Chromium Nuclei Isolation Kit followed by magnetic enrichment (n = 20 samples; sample-level details in Table S2), whereas the 10x Flex dataset comprised cerebellar samples prepared using sucrose pelleting followed by magnetic enrichment (n = 12 samples; run-level details in Table S1). Accordingly, this analysis is presented as an aggregate post-QC feasibility overview across the two sequencing workflows under the tested conditions, rather than as a controlled comparison of platform performance alone.

For this analysis, samples were first evaluated using a platform-aware QC framework incorporating run-level and sample-level metrics (Tables S1–S2). In the 10x Flex dataset, sequencing runs were flagged if Cell Ranger sequencing saturation was <20%, mean reads per cell was <15,000, or Q30 base quality was <85%, and were excluded if they failed the first two criteria. Using these thresholds, one multiplexed 10x Flex run (Set1) was excluded because it showed both low sequencing saturation (8.0%) and low mean reads per cell (9,725), while the remaining runs passed the hard exclusion criteria (Table S1). Probe-set mapping fraction and demultiplexing balance (Demux_pct) were monitored as informational run-level QC metrics; libraries with probe-mapping <70% or Demux_pct >50% were retained for sensitivity analyses but not used to define core performance.

In the PARSE WT dataset, sample-level QC focused on per-nucleus gene detection and mitochondrial content (Table S2). Samples were excluded if the 95th percentile mitochondrial read fraction exceeded 5%; accordingly, one sample with elevated mitochondrial signal (p95 mito = 8.28%; S12) was removed. Across both datasets, samples with fewer than 500 recovered nuclei were excluded outright, and samples with median detection below 1,000 genes per nucleus were flagged and retained only for sensitivity analyses in the pooled post-QC analysis (Table S2). Under these criteria, the remaining libraries and samples provided sufficient complexity and quality to support downstream cell-type–level and differential expression analyses.

To place these yields in context, reported gene detection in mouse brain snRNA-seq varies with sequencing depth and library preparation chemistry (platform/kit version), but droplet-based snRNA-seq can recover a few thousand genes per nucleus; for example, DroNc-seq reported an average of 2,731 genes per mouse brain nucleus when sequenced at ∼160,000 reads per nucleus (4). Separately, snRNA-seq has been successfully applied to the adult mouse hippocampal formation in large-scale atlas studies defining neuronal and non-neuronal cell types (13–15) and to the mouse cerebellum in atlas-scale datasets that define cerebellar cell types (16).

Across optimization experiments, workflows relying solely on OptiPrep gradients or Chromium Nuclei Isolation Kit-based isolation showed low recovery and substantial debris contamination in adult mouse myelin-rich brain tissue, often producing dilute suspensions (typically ≤10□nuclei/mL) with compromised morphology. In contrast, benchtop sucrose-gradient pelleting improved recovery and generated nuclei-dominant suspensions, routinely achieving ∼10□–10□nuclei/mL depending on tissue input and scaling. Addition of magnetic enrichment further reduced residual myelin/debris and improved purity with minimal additional loss of intact nuclei, yielding preparations compatible with downstream snRNA-seq library construction (1,3).

Tissue volumes for the stepwise yield schematic were estimated from mass using an assumed brain tissue density of 1.04 g/mL (17). Estimated total nuclei inputs were then approximated from literature-derived regional cell density values and cellular scaling relationships for mouse brain (18,19). These values were used only to provide a broad contextual framework for interpreting recovery and should not be interpreted as direct measurements of nuclei input for the samples analyzed here. Because regional cellularity can vary with biological and technical factors, including sex, age, strain, tissue composition, and sample handling, these estimates are inherently approximate. On this basis, hippocampus-scale inputs (∼20 mg) were estimated to contain on the order of ∼1–2 million nuclei, whereas cerebellum-scale inputs (∼63 mg; approximately half cerebellum) were estimated to contain ∼20–25 million nuclei. Relative to these approximate literature-based ranges, hippocampal workflows produced recoveries within a plausible operating range, whereas cerebellar samples showed greater losses during cleanup, consistent with the heavier debris burden of densely packed cerebellar tissue. These comparisons are intended to provide qualitative context rather than a precise quantitative benchmark of recovery efficiency (Fig. 5). Similarly, 10x Chromium kit–based workflows showed lower initial recovery but consistent post-cleanup retention (∼70–85% of pre-enrichment nuclei). In contrast, cerebellar preparations, despite achieving expected yields after sucrose pelleting (∼5–15% of total input), exhibited substantially lower recovery following cleanup/enrichment (∼10% of pre-enrichment nuclei) (Fig. 5). We attribute this reduced recovery primarily to the higher myelin/lipid content and fine debris burden of cerebellar tissue, as well as the smaller size and higher packing density of cerebellar granule cell nuclei, which increase susceptibility to loss during filtration, density separation, and cleanup steps. These observations highlight a tradeoff between purity and yield in myelin-rich brain regions, where more stringent debris removal improves sample cleanliness but can disproportionately reduce nuclei recovery, particularly at larger and more debris-rich inputs (1,3). A quantitative comparison of concentrations and yields across workflows is summarized in Table 1.

**Figure 5.**
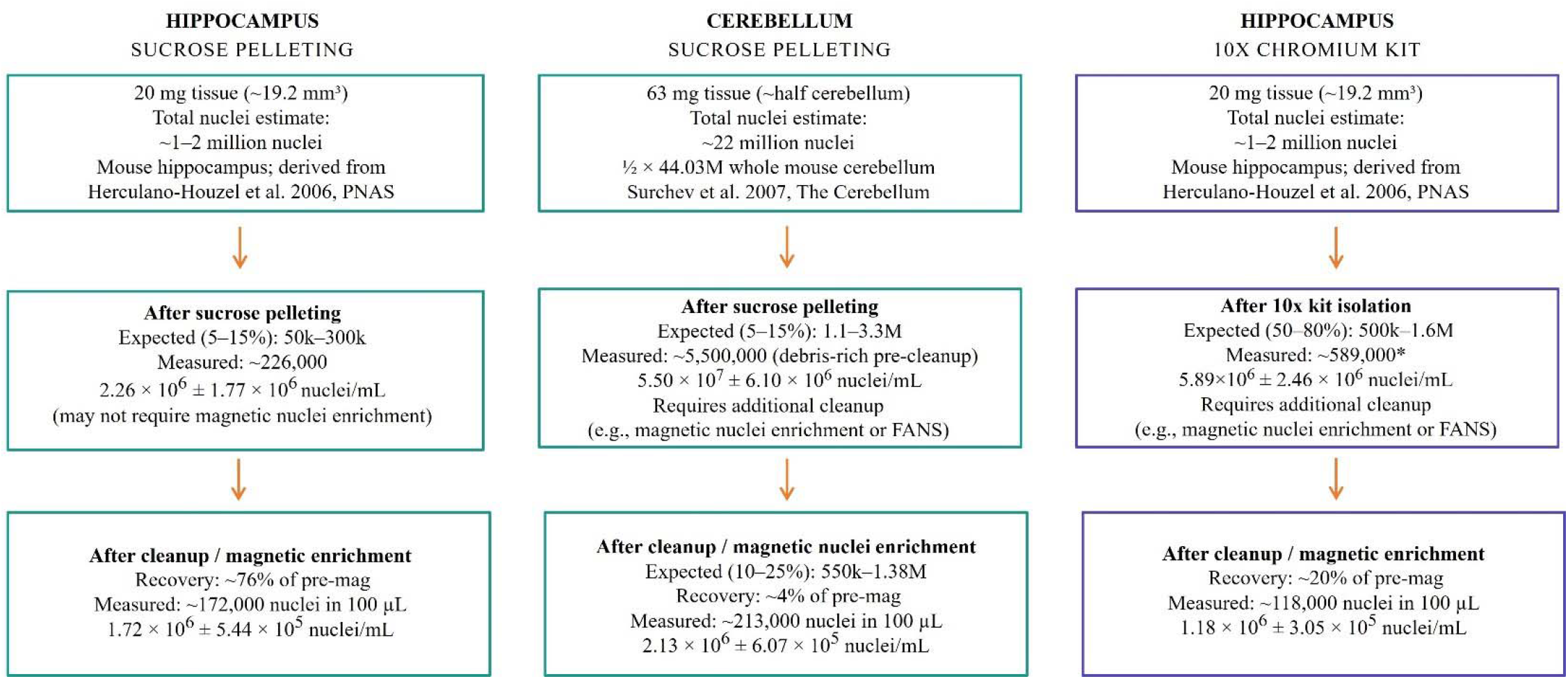
Estimated nuclei yields and stage-specific recovery for hippocampus and cerebellum workflows. Schematic showing expected and measured nuclei yields for hippocampus processed by sucrose pelleting (left), cerebellum processed by sucrose pelleting (middle), and hippocampus processed using the 10x Chromium Nuclei Isolation Kit plus magnetic enrichment (right). Total input nuclei were estimated from published regional cellularity references. For hippocampus (∼20 mg; ∼19.2 mm^3^), total input was estimated at ∼1–2 million nuclei. For cerebellum (∼63 mg; approximately half cerebellum), total input was estimated at ∼22 million nuclei. Boxes show expected yield ranges and measure nuclei counts after the initial isolation step and after cleanup/magnetic enrichment. In cerebellum, the post-pelleting count represents a debris-rich pre-cleanup preparation, and additional cleanup was required before final recovery was assessed. Overall, hippocampal workflows showed recovery within expected ranges, whereas cerebellar samples showed greater losses during cleanup, consistent with the heavier debris burden of densely packed cerebellar tissue. *For the 10x Chromium Kit hippocampal workflow, pre-enrichment counts include substantial debris that inflates automated nuclei counts; the low apparent recovery after cleanup primarily reflects debris removal rather than nuclei loss (see text, section Magnetic separation improves debris cleanup and Supplementary Figure 1)

**Table 1.**
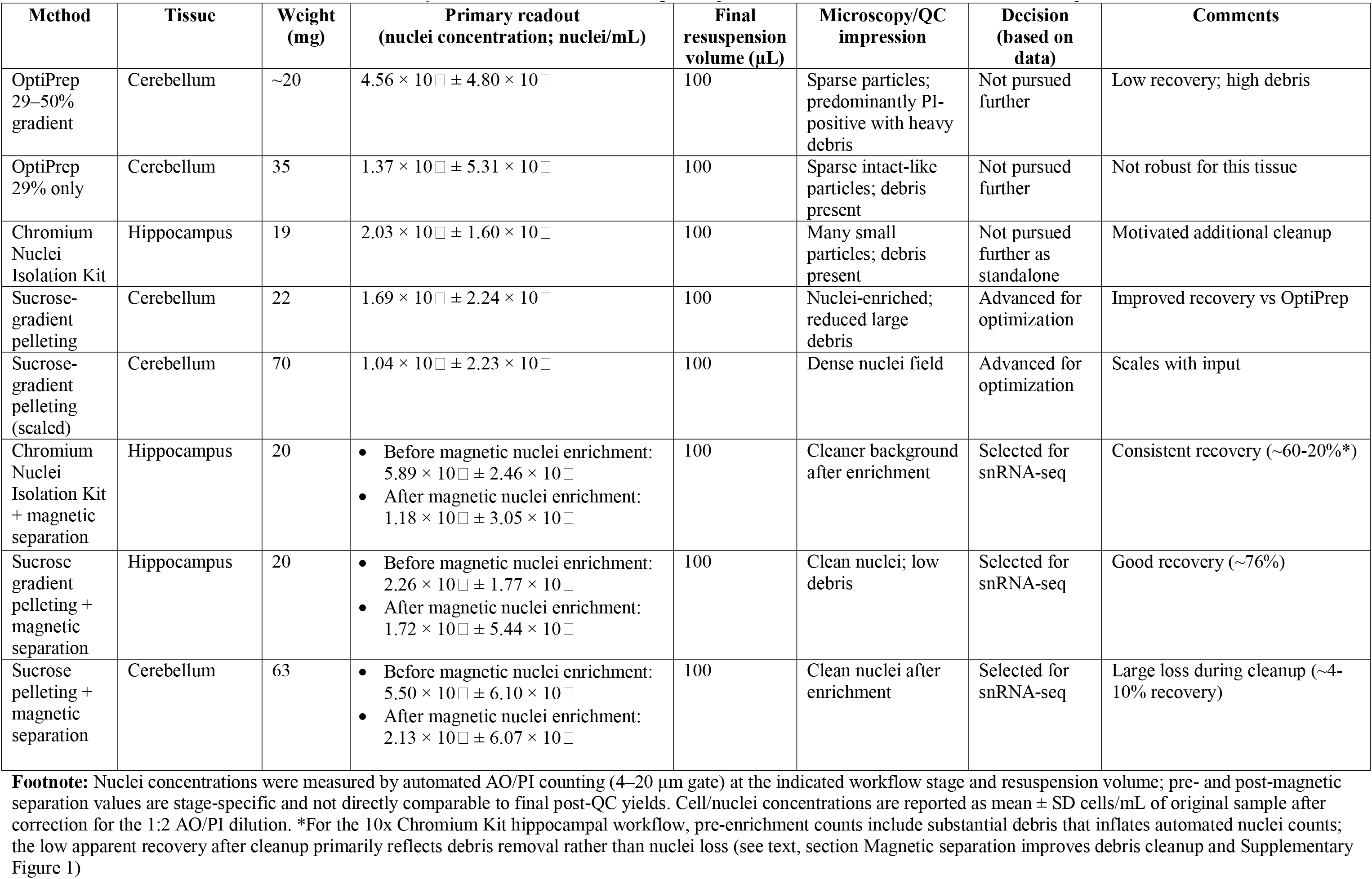
Summary of nuclei isolation and cleanup strategies evaluated for adult mouse brain snRNA-seq.

## LIMITATIONS

Nuclei isolation from adult, myelin-rich brain regions is highly sensitive to mechanical disruption and density-based cleanup conditions, and outcomes can vary substantially with brain region and biological context. Regions with high lipid/myelin content often require more stringent debris removal and nuclei integrity can further decline with increasing age, strain background, or disease states that alter myelination and tissue fragility (e.g., demyelination or neuroinflammation). Therefore, key parameters including homogenization intensity, detergent concentration, and cleanup stringency may require region- and context-specific optimization. This workflow is also vulnerable to sample handling losses during homogenization. Because homogenization is performed in small volumes and often produces foaming or splash-backs if executed too aggressively, minor sample spills can disproportionately reduce yield and introduce run-to-run variability. Minimizing spillage (e.g., keeping tubes on ice in a stable rack, using controlled strokes, and avoiding rapid plunging motions) is important for consistent recovery, particularly when tissue samples are limited.

In our hands, OptiPrep gradient conditions evaluated in adult mouse cerebella produced poor recovery and low-integrity nuclei, consistent with nuclei damage during handling and/or ineffective separation from lipid-rich debris. More broadly, OptiPrep-based workflows are less forgiving to small deviations in gradient preparation (e.g., concentration accuracy and layer stability), often require specialized consumables/equipment, and can become cost-limiting when scaled up due to reagent volume and waste. Conversely, benchtop sucrose pelleting was robust and scalable for myelin-rich adult tissue under our conditions but may still carry residual debris or rare cellular carryover depending on homogenization efficiency, filtration, and aspiration of the interphase; additional cleanup may be required for regions with high lipid content (5). Additionally, OptiPrep performance can be sensitive to gradient geometry and operator technique at the band-collection step. Widely used OptiPrep myelin-removal workflows are often optimized for larger-volume tubes (e.g., 15 mL) and ultracentrifugation, which can yield sharper interfaces and more reproducible banding. In this study, we adapted OptiPrep gradients to 2 mL tubes and a benchtop high-speed centrifuge, which may reduce band resolution and make recovery of the intermediate nuclei fraction more technically challenging, increasing the risk of layer mixing and nuclei loss. Under these equipment constraints, the benchtop sucrose pelleting approach reduced handling complexity and provided a more practical workflow for myelin-rich samples.

In this study, magnetic enrichment was treated as the recommended final cleanup step for sequencing-ready preparations from myelin-rich adult brain tissue, although some sucrose-pelleted preparations may appear sufficiently clean before this step. When omitted, users should verify low debris/myelin carryover by microscopy and counting before proceeding. In these contexts, magnetic enrichment can improve nuclei purity by reducing debris and carryover material, but it may also reduce apparent yield depending on column capacity, sample clumping, and losses during wash/elution. Importantly, magnetic cleanup improves purity but does not restore nuclei integrity if nuclei are already damaged upstream, underscoring the need to optimize homogenization and density separation prior to magnetic enrichment. For cerebellar samples, collecting the bottom pellet only after sucrose centrifugation may reduce debris carryover and improve starting purity, but this approach was not used as the default because it typically lowers nuclei recovery relative to collecting the pellet together with selected intermediate material. Tissue handling is an additional constraint as fresh versus frozen starting material can change nuclei integrity and debris profiles; freeze–thaw and prolonged storage may increase nuclear fragility and can elevate ambient RNA due to partial lysis. Even when the final suspension appears nuclei-enriched by microscopy, excessive shear or harsh lysis can release cytoplasmic RNA that increases “soup”/background in snRNA-seq and can distort cell-type signal. Users should therefore interpret preparations in conjunction with downstream QC metrics and, when possible, assess ambient RNA contributions using established computational approaches.

A further limitation of this study is that the PARSE WT libraries showed lower per-nucleus complexity than the 10x Flex libraries in the pooled post-QC comparison, with fewer genes detected per nucleus overall, while UMI counts were broadly comparable between platforms (Fig. 4). Cross-platform differences should be interpreted cautiously, as the number of genes and UMIs detected per nucleus can be influenced by library chemistry, sequencing depth, read retention, and sample-specific factors rather than reflecting a fixed platform-specific ceiling (20,21). In our dataset, the combination of adult, myelin-rich brain tissue and conservative sequencing depth for the PARSE subset likely contributed to reduced gene detection. Consequently, cross-platform differences in gene/UMI counts should be interpreted with caution when comparing absolute expression levels between technologies.

This protocol’s performance is also constrained by sample loading limits. Excess tissue mass or excessive suspension volume can reduce separation efficiency by collapsing or mixing the density interface, increasing debris carryover, and complicating pellet recovery. We did not design this study to quantitatively compare absolute nuclei yield across distinct brain regions; instead, our optimization emphasizes practical operating ranges in myelin-rich adult tissue. In particular, Chromium Nuclei Isolation Kit-based workflows may show effective performance only within a limited tissue-mass (in our experience, approximately 5–50 mg), whereas the sucrose benchtop method provides greater flexibility for input mass and total processing volume, especially when paired with optional magnetic cleanup for heavily myelinated samples.

Finally, this workflow is operator- and timing-dependent. Small variations in homogenization stroke number/speed, layering technique, aspiration depth, and total time on ice can meaningfully shift integrity and recovery. In addition, dye-based automated counter ‘viability’ readouts should be interpreted cautiously for nuclei: AO/PI (or similar) primarily reports permeability/dye access, not true viability, and some intact nuclei may be classified as ‘dead’ depending on staining conditions and instrument gating. For nuclei preparations, AO/PI readouts should be interpreted as nuclei (red) versus residual intact cells (green), rather than live versus dead cells. As a result, an apparently ‘low viability’ (high red fraction) sample can nevertheless be optimal for snRNA-seq, and microscopy plus downstream QC remain the most reliable indicators of preparation success.

### Future directions

More broadly, the field would benefit from standardized nuclei isolation protocols optimized for distinct brain cell types and contexts (e.g., region-, age-, and disease-specific adaptations), which may improve representation of fragile or low-abundance populations and enable more precise comparisons across studies and atlas-scale datasets.

## TROUBLESHOOTING

### Problem 1: Low nuclei integrity

**Possible causes:**Over-homogenization (excessive shear), tissue warming, harsh detergent exposure, prolonged processing time, or repeated harsh pipetting.

**Solutions:** Use fewer and slower homogenization strokes, keep all samples and reagents ice-cold, minimize pauses between steps, handle nuclei with wide-bore or low-retention tips, avoid vortexing, and use the minimum detergent concentration required for efficient nuclei release, adding detergent last with gentle mixing.

### Problem 2: Heavy myelin/debris carryover after density cleanup

**Possible causes:** Incomplete separation due to disturbed interface, insufficient pelleting time/RCF, improper aspiration of myelin cap/interphase, or overloaded input.

**Solution:** Layer samples slowly along the tube wall; avoid bubbles; confirm centrifuge settings in RCF (×g); remove the myelin cap first; aspirate the interphase slowly and leave a small residual volume above the pellet to prevent pellet loss; reduce input mass/volume if carryover persists; add optional magnetic cleanup downstream.

### Problem 3: Gradient collapses or layers mix during setup

**Possible causes:** Incorrect sucrose/OptiPrep concentration, temperature mismatch between layers, rapid pipetting, bubbles, or overfilling/overloading the tube.

**Solution:** Prepare gradients with accurate concentrations; pre-chill all solutions on ice; pipette slowly using the tube wall; avoid bubbles; do not exceed recommended sample volume; if collapse occurs, remake gradients and reduce loading volume.

### Problem 4: Too much sample volume causes poor separation and low recovery

**Possible cause:** Excess suspension volume reducing density contrast, increases mixing, and prevents efficient pelleting of intact nuclei.

**Solution:** Concentrate the sample before loading (gentle spin and resuspension in smaller volume); adhere to tube capacity and recommended layer volumes; process large inputs in parallel tubes rather than overloading a single gradient.

### Problem 5: Signs of incomplete homogenization (clumps, poor filtration, inconsistent counts)

**Possible causes:** Insufficient mechanical disruption, tissue not fully minced, or too few homogenization strokes. **Solution:** Mince tissue uniformly (∼1 mm³); increase strokes gradually using slow, controlled motion; visually confirm a uniformly turbid suspension without large fragments; if clogging occurs, perform an additional gentle homogenization step before filtration.

### Problem 6: Unexpected “contamination layers” or unclear phase boundaries after spin

**Possible causes:** High lipid content, incomplete myelin cap removal, mixed interface from poor layering, or residual intact cells.

**Solution:** Use consistent size gating (e.g., 4–20 µm) for counting; remove myelin cap carefully before aspirating the interphase; repeat wash + filtration; consider magnetic enrichment after pelleting for highly myelinated regions; if phases are unclear, reduce sample load and remake the density setup.

### Problem 7: Myelin layer is difficult to aspirate with a fine-bore tip (clogging, smearing, or disturbing the interface)

**Possible cause:** The myelin cap is viscous/adhesive and clogs narrow tips; direct aspiration can shear the cap and mix the interphase, increasing debris carryover and risking pellet loss.

### Solution

1. **Move the myelin cap aside rather than aspirating it first.**

`o` Using a pipette tip (or a wide-bore tip), gently push/slide the myelin sheet to the tube wall and “park” it there so it sticks (Fig. 6).
2. **Switch to a fresh tip** and remove the supernatant/interphase.

`o` Insert the tip down the side of the tube, keeping it away from the parked myelin layer.
`o` Aspirate the middle layer (debris-containing interphase) slowly.
3. **Protect the pellet.**

`o` Leave ∼100–200 µL above the bottom to avoid aspirating an often-invisible nuclei pellet.
4. **Alternative (if the cap must be removed):**

`o` Use a wide-bore tip to aspirate the myelin cap + interphase together in a controlled manner, then stop early and finish with a fresh tip near the bottom.

**CRITICAL:** The nuclei pellet may be small or not visible—aspirate slowly and stop early to prevent pellet loss.

**Figure 6.**
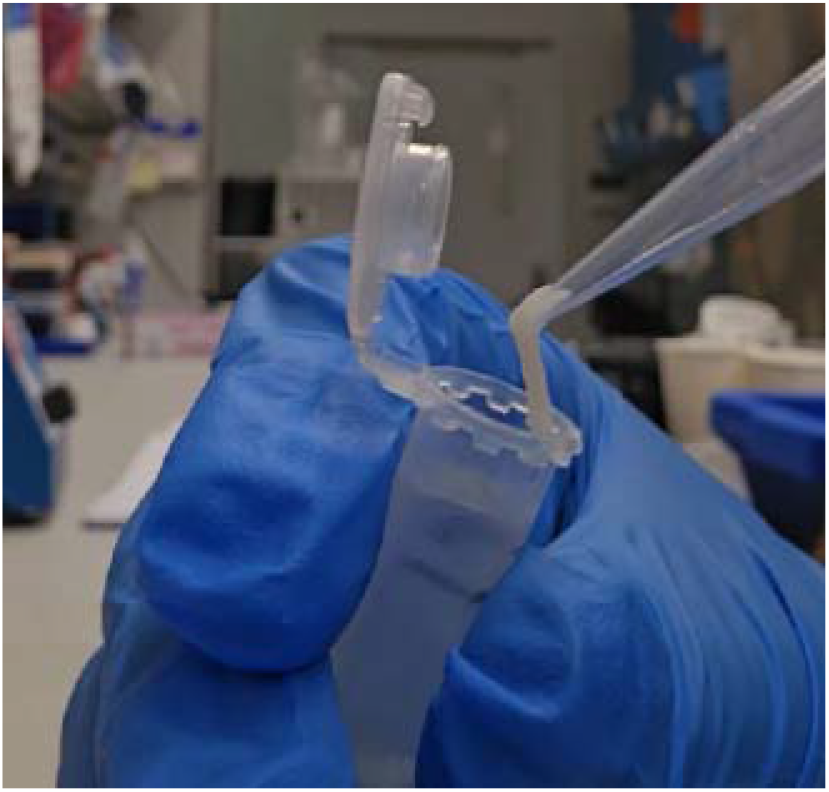
Technique for band collection to minimize layer disturbance during density-gradient cleanup. Representative example of aspirating an intermediate layer using a pipette tip positioned against the tube wall to reduce mixing of adjacent gradient phases and minimize nuclei loss during band recovery.

### Problem 8: Pellet is not visible, or nuclei are lost during aspiration

**Possible causes:** Nuclei pellet may be small/invisible; aspiration too aggressive; pellet dislodged.

**Solution:** Assume the pellet is present even if not visible; aspirate slowly; leave ∼100–200 µL above the pellet; mark tube orientation; use a narrow tip only for the upper layers and switch to wide-bore near the bottom; resuspend gently.

### Problem 9: Automated counter disagreement (e.g., Denovix vs Luna) or inconsistent nuclei numbers

**Possible causes:** Differences in gating/classification, dye chemistry timing, mixing consistency, or debris misclassification.

**Solution:** Standardize staining incubation time and mixing; use the same size gate across experiments; validate with microscopy; if possible, calibrate against a hemocytometer for a subset of samples; prioritize downstream snRNA-seq QC as the final validation.

## IACUC APPROVAL

All mouse procedures were approved by the University of Iowa Institutional Animal Care and Use Committee (IACUC protocol no. 4082613) and performed in accordance with institutional guidelines and the *Guide for the Care and Use of Laboratory Animals*.

## ACKNOWLEDGMENTS

We gratefully acknowledge Dr. Nandakumar Narayanan for critical reading of the manuscript.

## FUNDING SOURCES

National Institutes of Health grant R03 AG078787 (QZ) National Institutes of Health grant R01 AG086396 (QZ)

## AUTHOR CONTRIBUTIONS

BG and QZ designed the experiments. BG and BK performed all experiments. BG and QZ performed all data analyses. BG wrote the manuscript, BK and QZ reviewed and revised the manuscript.

## CONFLICTS

The authors declare that there are no conflicts of interest.

## Notes

**Conflict of Interest:** There are no conflicts of interest.

### Competing Interest Statement

The authors have declared no competing interest.

### Summary of Updates

The manuscript was revised to add Victor Kilonzo to the author list in recognition of his contributions to the project. Figure 3 was added, and the related data, results, figure legend, and reported recovery values were updated for consistency. The protocol text was also revised to improve clarity and reproducibility, including clearer instructions for sucrose cushion preparation, sample layering, resuspension volumes, and removal of residual sucrose. A supplementary file was also added. Minor grammatical and formatting corrections were made. The overall scope and conclusions of the manuscript remain unchanged.

