## Supplementary data for "Myelin-Free Nuclei Isolation from Mouse Hippocampus and Cerebellum for snRNA-Seq with Benchtop Gradient Centrifugation"

Sample information

a 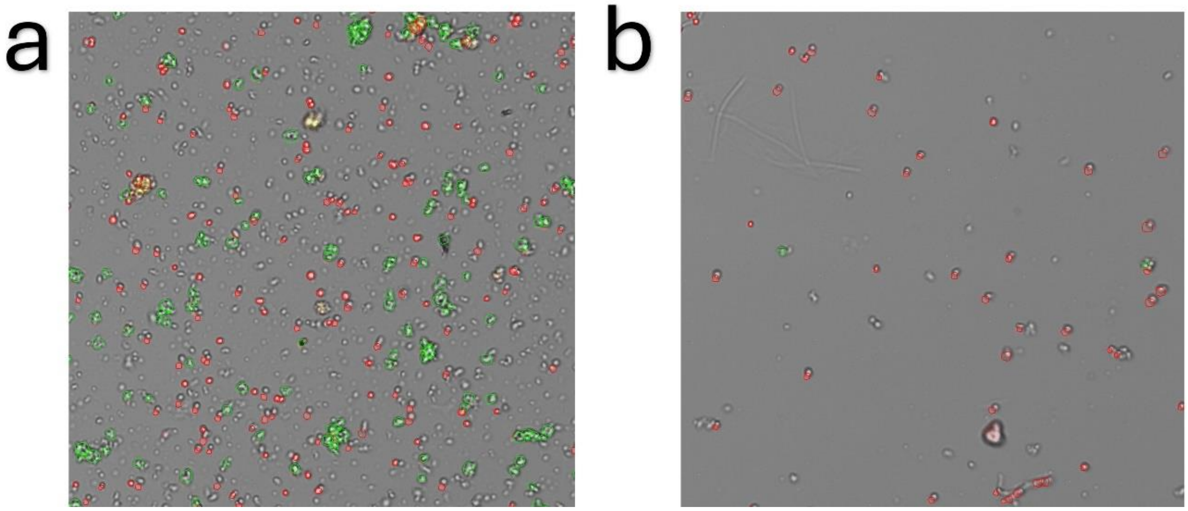

Supplementary Figure 1. **Representative images of hippocampal nuclei preparations before magnetic cleanup.**

(a) Sucrose pelleting–based hippocampal preparation after sucrose pelleting alone. The suspension is largely myelin- and debris-free, with predominantly intact nuclei visible, illustrating that additional magnetic cleanup is optional in this context and was used primarily to quantify potential losses. (b) Chromium Nuclei Isolation Kit–based hippocampal preparation prior to magnetic enrichment. Abundant debris and small particles are present and are frequently classified as “nuclei” by the automated counter, leading to overestimation of pre-enrichment nuclei counts. These images illustrate the different debris burdens underlying the yield and “recovery” metrics reported in the main text.

**Table S1:** Sequencing run-level QC metrics and pooling design for 10x Genomics multiplexed snRNA-seq libraries.

| Sample label | Saturation_pct | Mean_reads_per_cell | Reads_in_cells_pct | Probe_mapping_pct | Q30_rna_pct | Demux_pct | QC flag(s) | Final decision |
|---|---|---|---|---|---|---|---|---|
| Set1_5-1 | 8.0 | 9725 | 83.7 | 94.3 | 92.4 | 24.6 | Saturation < 20%; Mean reads per cell < 15,000 | Excluded |
| Set1_6-1 | 8.0 | 9725 | 83.7 | 94.3 | 92.4 | 17.5 | Saturation < 20%; Mean reads per cell < 15,000 | Excluded |
| Set1_7-1 | 8.0 | 9725 | 83.7 | 94.3 | 92.4 | 27.2 | Saturation < 20%; Mean reads per cell < 15,000 | Excluded |
| Set1_8-1 | 8.0 | 9725 | 83.7 | 94.3 | 92.4 | 30.7 | Saturation < 20%; Mean reads per cell < 15,000 | Excluded |
| Set2_5-2 | 47.1 | 24469 | 82.3 | 92.8 | 87.2 | 5.7 | None | Included |
| Set2_6-2 | 47.1 | 24469 | 82.3 | 92.8 | 87.2 | 6.2 | None | Included |

| | | | | | | | | |
|---|---|---|---|---|---|---|---|---|
| Set2_7-2 | 47.1 | 24469 | 82.3 | 92.8 | 87.2 | 25.1 | None | Included |
| Set2_8-2 | 47.1 | 24469 | 82.3 | 92.8 | 87.2 | 63.0 | Demux_pct > 50% | Included (sensitivity) |
| Set3_5-4 | 82.8 | 38572 | 82.5 | 80.7 | 90.1 | 5.9 | None | Included |
| Set3_6-4 | 82.8 | 38572 | 82.5 | 80.7 | 90.1 | 8.8 | None | Included |
| Set3_7-4 | 82.8 | 38572 | 82.5 | 80.7 | 90.1 | 51.6 | Demux_pct > 50% | Included (sensitivity) |
| Set3_8-3 | 82.8 | 38572 | 82.5 | 80.7 | 90.1 | 33.7 | None | Included |
| Set4_5-5 | 79.6 | 36598 | 77.6 | 68.1 | 88.7 | 15.4 | Probe_mapping_pct < 70% | Included (sensitivity) |
| Set4_6-5 | 79.6 | 36598 | 77.6 | 68.1 | 88.7 | 12.2 | Probe_mapping_pct < 70% | Included |

| | | | | | | | | |
|---|---|---|---|---|---|---|---|---|
| | | | | | | | | (sensitivity) |
| Set4_7-5 | 79.6 | 36598 | 77.6 | 68.1 | 88.7 | 34.6 | Probe_mapping_pct < 70% | Included (sensitivity) |
| Set4_8-4 | 79.6 | 36598 | 77.6 | 68.1 | 88.7 | 37.7 | Probe_mapping_pct < 70% | Included (sensitivity) |
| Libraries were generated in four multiplexed pools. Pool 1 contained female wild-type mice treated with Drug (libraries Set1_5-1, Set2_5-2, Set3_5-4, Set4_5-5). Pool 2 contained male wild-type mice given Water (Set1_6-1, Set2_6-2, Set3_6-4, Set4_6-5). Pool 3 contained female hemizygous mice given Water (Set1_7-1, Set2_7-2, Set3_7-4, Set4_7-5). Pool 4 contained male hemizygous mice treated with Drug (Set1_8-1, Set2_8-2, Set3_8-3, Set4_8-4). | | | | | | | | |

**Table S2:** Sample-level mouse metadata and snRNA-seq nuclei QC metrics for PARSE WT dataset

| **Sample** | **Condition** | **Strain / genotype** | **Sex** | **n_cells** | **Median genes** | **Median UMIs** | **Median mito** | **p95 mito** | **QC flag(s)** | **Final decision** |
|---|---|---|---|---|---|---|---|---|---|---|
| S1 | Calorie restricted | C57BL/6 | Male | 1721 | 2068 | 5387 | 0 | 0.055556 | None | Included |
| S2 | Control | C57BL/6 | Male | 773 | 2520 | 7713 | 0 | 0.021672 | n_cells < 800 | Included (sensitivity analysis) |

| | | | | | | | | | | |
|---|---|---|---|---|---|---|---|---|---|---|
| S3 | Aged | C57BL/6 | Male | 1935 | 1893 | 4732 | 0 | 0.032782 | None | Included |
| S4 | Water | Wild-type littermate | Male | 4216 | 2150.5 | 5187 | 0.007386 | 0.181769 | None | Included |
| S5 | Drug | 5xFAD | Male | 2132 | 2114.5 | 5362.5 | 0.01974 | 0.265456 | None | Included |
| S6 | Drug | 5xFAD | Male | 3097 | 1769 | 4449 | 0 | 0.071713 | None | Included |
| S7 | Drug | Wild-type littermate | Male | 572 | 1448 | 2913 | 0 | 0.108203 | n_cells < 800 | Included (sensitivity analysis) |
| S8 | Drug | Wild-type littermate | Male | 6105 | 991 | 1848 | 0.021626 | 0.30448 | Median genes < 1000 | Included (sensitivity analysis) |
| S9 | Water | 5xFAD | Male | 2645 | 959 | 1684 | 0 | 0.240743 | Median genes < 1000 | Included (sensitivity analysis) |
| S10 | Water | 5xFAD | Male | 3294 | 831 | 1410.5 | 0.008092 | 0.341706 | Median genes < 1000 | Included (sensitivity analysis) |
| S11 | Water | Wild-type littermate | Male | 4795 | 740 | 1237 | 0.054166 | 0.559887 | Median genes < 1000 | Included (sensitivity analysis) |
| S12 | Control | C57BL/6 | Male | 531 | 2284 | 5912 | 0.030957 | 8.281927 | n_cells < 800; p95 | Excluded |

| | | | | | | | | | | |
|---|---|---|---|---|---|---|---|---|---|---|
| | | | | | | | | | mito > 0.05 | |
| S13 | AAV-knockdown model | C57BL/6 | Male | 2036 | 1743.5 | 4044 | 0.036812 | 2.484634 | None | Included |
| S14 | Control | C57BL/6 | Male | 1660 | 1676.5 | 3940.5 | 0.006939 | 0.119175 | None | Included |
| S15 | AAV-knockdown model | C57BL/6 | Male | 4219 | 1494 | 3303 | 0 | 0.136908 | None | Included |
| S16 | Control | C57BL/6 | Female | 7117 | 1343 | 2958 | 0 | 0.088433 | None | Included |
| S17 | AAV-knockdown model | C57BL/6 | Female | 4478 | 1618.5 | 3738.5 | 0 | 0.106696 | None | Included |
| S18* | Control | C57BL/6 | Female | 8535 | 1511 | 3367 | 0 | 0.089298 | None | Included |
| S19 | AAV-knockdown model | C57BL/6 | Female | 8608 | 1566 | 3661 | 0 | 0.095287 | None | Included |
| S21 | Control | C57BL/6 | Female | 2990 | 1150 | 2266 | 0 | 0.133489 | None | Included |
| S22 | AAV-knockdown model | C57BL/6 | Female | 3611 | 1338 | 2888 | 0 | 0.102934 | None | Included |
